# Drivers of taxonomic, functional and phylogenetic diversities in dominant ground-dwelling arthropods of coastal heathlands

**DOI:** 10.1101/2021.01.04.425176

**Authors:** Axel Hacala, Denis Lafage, Andreas Prinzing, Jérôme Sawtschuk, Julien Pétillon

## Abstract

Although functional and phylogenetic diversities are increasingly used in ecology for a large variety of purposes, their relationships remain unclear and likely vary presumably over taxa, yet most recent studies focused on plant communities. Different concepts predict that a community becomes functionally more diverse by adding phylogenetic lineages, subtracting lineages, adding species, reducing or increasing environmental constraints. In this study, we investigated ground-dwelling spider, ground beetle and ant assemblages in coastal heathlands (>11 000 individuals, 216 species), and their estimated functional and phylogenetic diversities as minimum spanning trees using several traits related to the morphology, feeding habits and dispersal of species, and phylogenetic trees, respectively. Correlations were overall positive and high between functional and phylogenetic diversities. Accounting for taxonomic diversities and environments made disappear this relationship in ants, but maintained them in spiders and ground beetles, where taxonomic diversity related to functional diversity only via increasing phylogenetic diversity. Environmental constraints reduced functional diversity in ants, but affected functional diversity only indirectly via phylogenetic diversity (ground beetles) and taxonomic and then phylogenetic diversity (spiders and ground beetles). Results are consistent with phylogenetic conservatism in traits in spiders and ground beetles, while in ants traits appear more neutral with any new species potentially representing a new trait state. Lineage diversities mostly increased with taxonomic diversities, possibly reflecting un-measured environmental conditions.

## Introduction

Growing in popularity during the last twenty years (Webb et al. 2002; Campbell et al. 2010), functional and phylogenetic metrics have been very useful in a large variety of contexts. In applied ecology for instance, phylogenetic diversity (PD) was successfully used to establish conservation prioritization (Rodrigues et al. 2005; Magura 2016; Tucker et al. 2018; Wong et al. 2019). PD was also proven relevant for understanding community assembly rules and underlying interactions (Cavender-Bares et al. 2009). The same is true for functional diversity (FD), with cases where functional diversity led to a better understanding of ecological processes that classical taxonomic diversity alone did not properly assess (Leroy et al. 2014; Campbell et al. 2010; Wong et al. 2019). There are different approaches to quantifying PD and FD (Pavoine et al. 2013). We here stick to the approach particularly appropriate in a conservation context, i.e. measuring PD and FD represented by the sum of all species present, such as the size of a minimum spanning tree connecting all species in phylogenetic of functional space (Faith 1992).

While the link between the phylogeny and traits is considered strong due to phylogenetic conservatism (Darwin 1859 in Webb et al. 2002, Wiens 2010), this conservatism has very rarely been tested locally (Pavoine et al. 2010) and hence the links between local PD and FD are not fully elucidated (Cadotte et al. 2019). Only if local phylogenetic conservatism is perfect does an increase in PD inevitably increase FD (Webb et al. 2002). However, locally, phylogenetic conservatism may not be perfect or even be absent (Eterovick et al. 2010, Haak et al. 2014, Marcillo-Silva et al. 2016, Grundler et al. 2018). In that case, other patterns might theoretically emerge. First, FD might be unrelated to PD. In the most simple, neutral, model of community assembly (Hubbell 2001), traits are unimportant and any species arriving or going extinct carries a random chance of representing a unique trait state within a given community (Purves & Turnbull 2010, Novack-Gottshall 2016). FD would increase with taxonomic diversity (TD), not with PD. In niche-based models, particular environmental constraints might select against certain functional trait states and hence reduce FD (Webb et al. 2002), but might also impose character displacement among species and reduce niche packing (Safi et al. 2011) and thereby increase FD. Finally, character displacement may occur among competing close relatives leading to a negative rather than a positive relationship between PD and FD (Prinzing et al. 2008).

Despite this diversity of theoretically possible relationships between PD and FD, several authors argued that PD is a proxy of FD, and that the more traits are used to calculate FD the more it will resemble PD (Flynn et al. 2011; Khalil et al. 2018; Tucker et al. 2018). As recently highlighted by Cadotte et al. (2019), most studies showed a positive correlation between phylogenetic and functional diversity (PFC). Yet the spectrum of correlation strength is wide, with a few studies even reporting negative correlations (Bernard-Verdier et al. 2013). This gray area led to debate about the relationships between PD and FD (Pavoine et al. 2013; Tucker et al. 2018), and highlights the need for more empirical studies. Yet, most of previous work was done on plant communities (Cavender-Bares et al. 2009; Campbell et al. 2010), this redundancy of study models being recurrent in ecology, and easily fixed by multiplying model taxa (Coelho et al. 2009; Gerlash et al. 2013;Wong et al. 2019).

Our study therefore proposes to compare the PFC between several dominant taxa of arthropods using a standardized sampling protocol taking advantage of a large-scale restoration design on maritime cliff-tops, and to assess whether taxonomic diversities and/or abiotic forces drive both functional and phylogenetic diversities in each taxa. We studied three groups of arthropods, spiders, ground beetles and ants. These arthropods are known to complement vegetation survey (Pétillon et al. 2014; Hacala et al. 2020), and thereby offer an interesting opportunity to better understand the link between PD and FD (Wong et al. 2019). Spiders, ants and ground beetles are dominant ground-active macro-arthropods in several temperate habitats, and their traits and phylogeny are relatively well known (Bond et al. 2014; Pedley et al. 2014; Schirmel et al. 2016; Magura 2016; Parr et al. 2017). Up to our knowledge, one publications has already treated PD and FD for a single group of ground dwelling arthropods (Arnan et al. 2015), and some have conducted simple bivariate correlations between FD to PD (Corbelli et al. 2015; Liu et al. 2016), all having observed a positive PFC (but see Ridel et al. subm.). The novelty of our study resides in the comparison between several ground dwelling taxa, and in the fact that we look for more than just bivariate correlation but investigate the complex driving forces behind patterns of taxonomic functional and phylogenetic diversities of spiders, carabids and ants. In this study, we expect 1) environmental drivers of FD and PD to differ between taxa, reflecting different habitat filtering and 2) similar phylogenetic constraints in sympatric taxa of arthropods.

## Methods

### Sampling and identification

Sampling took place in three coastal sites of Brittany, Western France in 2017: L’Apothicairerie (47° 21′ 44.0″ N, 3° 15′ 34.9″ W), La Pointe de l’Enfer (47° 37′ 18.3″ N 3° 27′ 46.9″ W) and La Pointe de Pen-Hir, located on the mainland (48° 15′ 03″ N, 4° 37′ 25″ W). These sites were selected for they are all comparable with similar dominant vegetation (a short and dry heathland dominated by *Erica spp*. and *Ulex spp*), and under ongoing ecological passive restoration with 3 degradation states by site (see full description of study sites and pictures in Hacala et al. 2020).

Two 400 m^2^ plots of homogeneous vegetation were set for each degradation state, and four pitfall traps (80 mm in diameters and 100 mm deep) were set at each plot. Traps were half-filled with a salted solution (250 g L^−1^) with a drop of odorless soap and settled 10 meters apart in order to avoid interference and local pseudoreplication (Topping and Sunderland 1992). This resulted in 71 traps (in one station, the sampling area was too restricted to set 4 traps spaced of 10 m apart, so 1 was removed) active between mid-March to mid-June 2017, and emptied every 2 weeks. Total plant cover was estimated in a 5 m radius circle around each trap, and all species were identified and their percentage cover estimated. Environmental variables where inferred from the vegetation communities using Ellenberg’s index values (1992) extracted from Hill *et al*. (2004) and corrected for the British Isles. Anthropogenic degradation was assessed by using bare ground as a proxy of degradation intensity (Hacala et al. 2020). Pitfall samples were sorted in laboratory, arthropods transferred to ethanol 70°, and stored at the University of Rennes 1. Spiders, grounds beetles and ants were identified to species level. Spiders were identified using Roberts (1985) and Nentwig et al. (2019). Ground beetles were identified using Luff (2007) and Jeannel (1941). Ants were identified to species levels using Blatrix et al. (2013).

### Trait gathering

Three criteria were set to select the traits for calculating functional diversity of all three taxa. (1) They must be available in the literature for the three taxa. (2) They must be reported as drivers of species assemblages for each taxon. (3) They exist under a form or can be transformed in a way to be comparable between taxa (*e*.*g*. numerical into categorical). The functional trait definition provided by Violle et al. (2007) and followed by Wong et al. (2019) was finally used, with the exclusion of environmental preferences as they are not strictly speaking a functional trait. Three trait were selected and their values gathered from existing literature: dispersal abilities, body size and trophic guild (Table 1), all these traits being known to be affected by environmental responses in spider, carabid and ant assemblages (Schirmel et al. 2010; Lafage et al. 2015). Dispersal trait was chosen to express dispersal at a landscape scale in order to account for the colonization ability of the studied species. The spider capacity to disperse by the air (ballooning) was the criterion used to discriminate dispersal ability. For ground beetles, the ability to fly based on wing development was used for dispersal ability. For the ants, the queen’s way of locomotion for founding a new colony (by flight or by foot after a colony division) was used as comparable landscape level of dispersal. Body size was divided into three discreet classes to facilitate the comparison with the other variables. The division into size classes was done for each class to contain a comparable number of species. The guild trait was acquired through the feeding regime for ants and ground beetles. For spiders, the foraging strategy was used to distinguish guilds inside a predator-only taxon. We stress that traits were never inferred from phylogenetic position (see Table 1 for details and literature used).

**Table 1:**
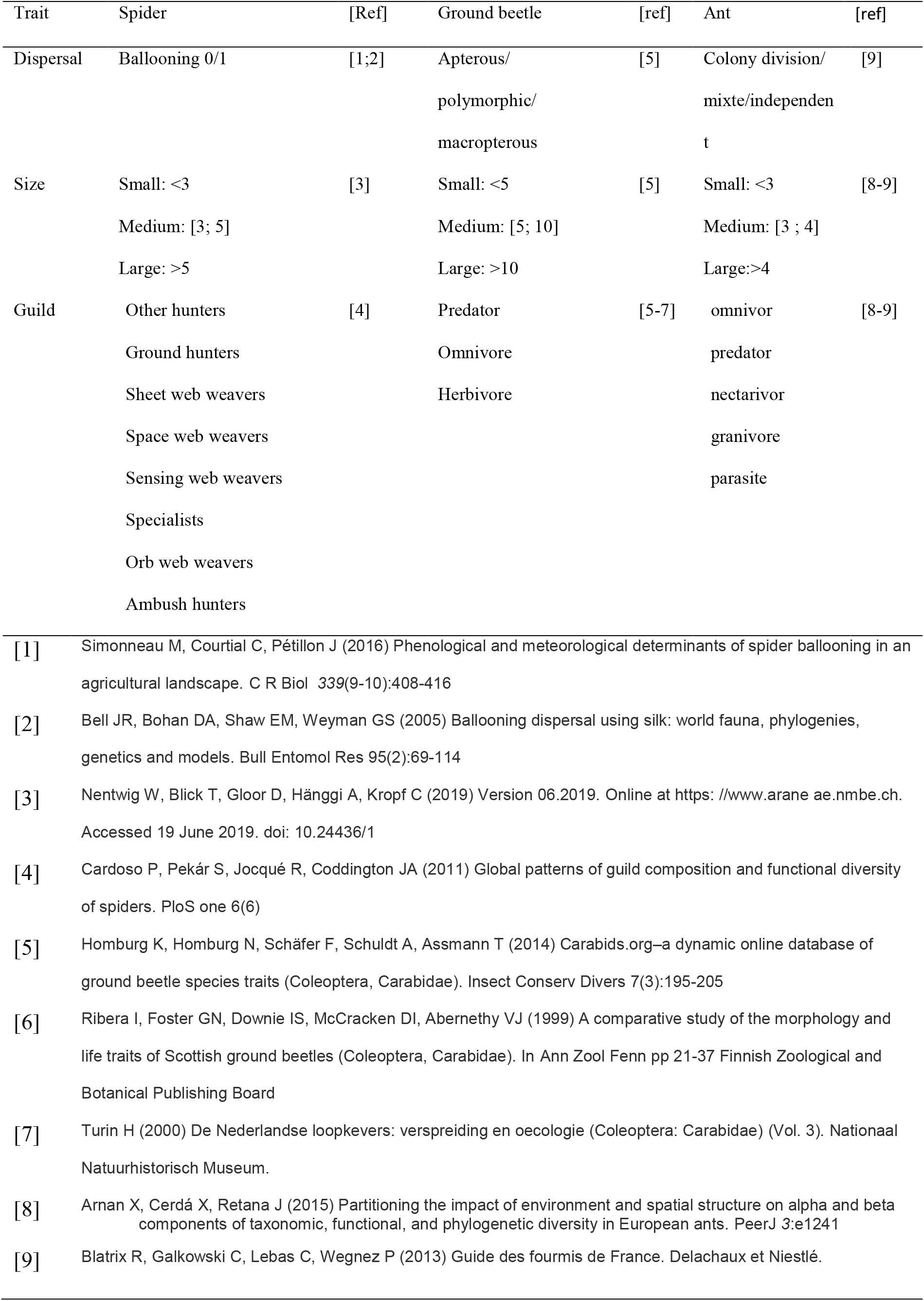
Modalities of functional trait for each taxa.

### Phylogenetic tree building

Phylogenetic trees were constructed by combining phylogenetic and taxonomic data from literature, assuming identical branch length between genus (1) and species (0.5) as real branch lengths because sequences were not available for all identified species.

The spider phylogenetic tree was adapted from Wheeler et al. (2017). Genera that were not present in Wheeler’s tree were placed using Arnedo et al. (2009), Frick et al. (2010) and Wang et al. (2015) for Linyphiidae and Agnarsson (2004), Azevedo et al. (2018), Maddison (2015), Millidge (1977), Piacentini & Ramírez (2019) and Scharff et al. (2019) for other families. The carabid phylogenetic tree was adapted from López-López & Vogler (2017), Martínez-Navarro et al. (2005), Ober & Maddison (2008), Ruiz et al. (2009) and Sasakawa & Kubota (2007). The ant phylogenetic tree was adapted from Moreau et al. (2006)

### Diversity calculating

The two diversity metrics, PD and FD, were calculated using the BAT package (Cardoso et al. 2015). Distance matrices were computed with gower distance from the FD package (Laliberté et al. 2014) for functional distances and with as.phylo from the ape() package (Paradis et al. 2019) for phylogenetic distances. PD and FD were calculated by meaning the Jackknifes estimates from apha.estimated () function from BAT in order to account for sampling variability.

### Statistical analysis

Correlations between PD and FD were estimated in a Bayesian framework with a Student’s t distribution with the brms package (Bürkner 2018). We used 2000 iterations on 4 chains. Model convergence was checked by visually inspecting diagnostic plots.

To select environmental variables affecting PD and FD, models were built within a Bayesian framework using brms (Bürkner, 2018) with two chains and default priors. All environmental variables were standardized and centered. The models included % bare-ground, litter depth, soil depth, and vegetation height. It also included community-weighted means of nitrogen level, light, salinity, pH and humidity based on Elenberg indicator values (modified by Hill et al., 1999). Model convergence was checked by visually inspecting diagnostic plots and using Rhat value. Parameter selection was based on “HDI+ROPE decision rule” (Kruschke & Liddell 2018) with a range value determined as −0.1 * sd(y), 0.1 * sd(y) (Kruschke & Liddell 2018) and was performed using bayestestR (Makowski et al., 2019).

We assessed the relative contribution of environmental variables selected by Bayesian models using structural equation modeling (SEM). The SEM approach also allowed us to assess the links between taxonomic diversity (TD), PD and FD taking environment into account. A significant correlated error between the two variables would indicate the existence of an unknown parameter influencing both variables. We used the piecewiseSEM package (Lefcheck 2016) as it allows using mixed models in association with nlme package (Pinheiro et al. 2020). Our initial model included the following links: (1) PD is affected by TD and selected environmental variables, (2) FD is affected by TD, selected environmental variables and PD, (3) TD is affected by selected environmental variables and (4) there is correlated error between PD and FD. Site was used as a random factor in every link modeled using nlme (Pinheiro et al. 2019). After the specification of the initial model, we re-defined our model excluding non-significant links (p<0.05) using a stepwise approach until ΔAICc <2 between two subsequent models. Finally, we assessed model fit using Fisher’s C statistic.

## Results

### Simple correlations

Correlations between phylogenetic and functional diversities were 0.59 (95% CI: 0.41-0.73), 0.65 (95% CI: 0.46-0.79) and 0.72 (95% CI: 0.57-0.82) for spiders, carabid beetles and ants respectively (Fig. 1), and overall increased with decreasing species richness of taxa (153, 40 and 23, respectively).

**Figure 1:**
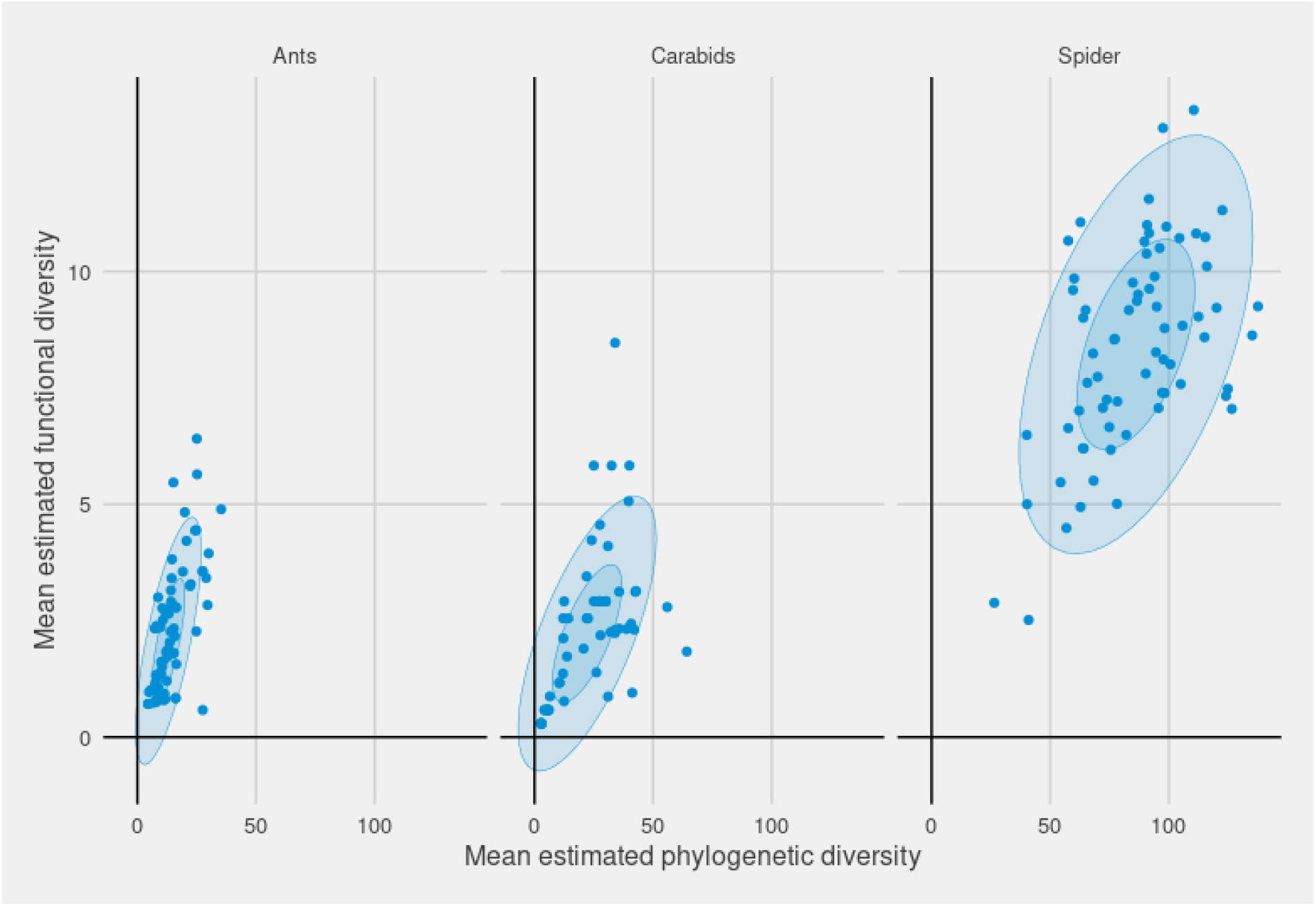
Plot of mean estimated functional diversity as a function of phylogenetic diversity. Blue ellipses correspond to 95% and 5% normal confidence ellipses. R^2^ = 0.384; 0.423, and 0.518. Se Figure 2 for a broader picture accounting also for taxonomic diversity and the environment.

### Variable pre-sélection

The model for spider phylogenetic diversity successfully converged and had a R^2^ = 0.589. Humidity and salinity were the best explanatory variables. Humidity effect on spider phylogenetic diversity had a high probability of existing (pd = 99.6%, Median = 16.37, 89% CI [7.45, 26.31]) and could be considered as significant (0% in ROPE). Salinity effect on spider functional diversity had a high probability of existing (pd = 97.8%, Median = −21.00, 89% CI [-38.68, −4.90]) and could be considered as significant (0% in ROPE). The model for spider functional diversity successfully converged and had a R^2^ = 0.26, nevertheless, none of the environmental variables could be considered as significant (ROPE >0%). The model for spider TD successfully converged and had a R^2^ = 0.66. Humidity and salinity were the best explanatory variables. Humidity effect on spider TD had a high probability of existing (pd = 99.4%, Median = 0.56, 89% CI [0.27, 0.92]) and could be considered as significant (0% in ROPE). Salinity effect on spider TD had a high probability of existing (pd = 99.4%, Median = −0.98, 89% CI [-1.56, −0.34]) and could be considered as significant (0% in ROPE).

The model for carabid beetle phylogenetic diversity successfully converged and had a R^2^ = 0.44. Percent bare-ground and salinity were the best explanatory variables. Percent bare ground effect on carabid phylogenetic diversity had a high probability of existing (pd = 99.2%, Median = −6.90, 89% CI [-11.73, −2.67]) and could be considered significant (0% in ROPE). Salinity effect on carabid phylogenetic diversity had a high probability of existing (pd = 99.9%, Median = 21.01, 89% CI [8.11, 32.19]) and could be considered significant (0% in ROPE). The model for carabid functional diversity successfully converged and but had a low R^2^ = 0.24. Salinity was the best explanatory variable. Salinity effect on carabid functional diversity had a high probability of existing (pd=99.4%, Median = 2.61, 89%CI [0.78, 4.09]) and could be considered significant (0% in ROPE). The model for carabid TD successfully converged with R^2^ = 0.44. Salinity was the best explanatory variable. Salinity effect on carabid TD had a 89.2% probability of existing (Median = 0.64, 89%CI [-0.29, 1.45]) and could be considered significant (0% in ROPE)

The model for ant phylogenetic diversity successfully converged and had a R^2^ = 0.31. Soil depth was the best explanatory variable. Soil depth effect on spider phylogenetic diversity had a high probability of existing (pd = 97.5%, Median = 2.84, 89% CI [0.68, 5.11]) and could be considered as significant (0.4% in ROPE). The model for ant functional diversity successfully converged and had a R^2^ = 0.40. Soil depth was the best explanatory variable. Soil depth effect on ant functional diversity had a high probability of existing (pd = 99.10%, Median = 0.59, 89% CI [0.18, 0.99]) and could be considered as significant (0% in ROPE). The model for ants TD successfully converged and had a R^2^ = 0.37. Soil depth was the best explanatory variable. Soil depth effect on ant TD had a medium probability of existing (pd = 98.10%, Median = 0.36, 89% CI [0.09, 0.66]) and could be considered as significant (1.1% in ROPE).

### Structural equation models

When testing the relationship between diversity metrics and environmental variables, our final SEMs indicated good fit with the data both for spiders (Fisher’s C = 17.19, p = 0.07; Fig. 2a) and carabid beetles (Fisher’s C = 9.77, p = 0.636, Fig. 2b). The fit of the ant model could not be estimated as the best model was fully saturated (Fig. 2c).

**Figure 2:**
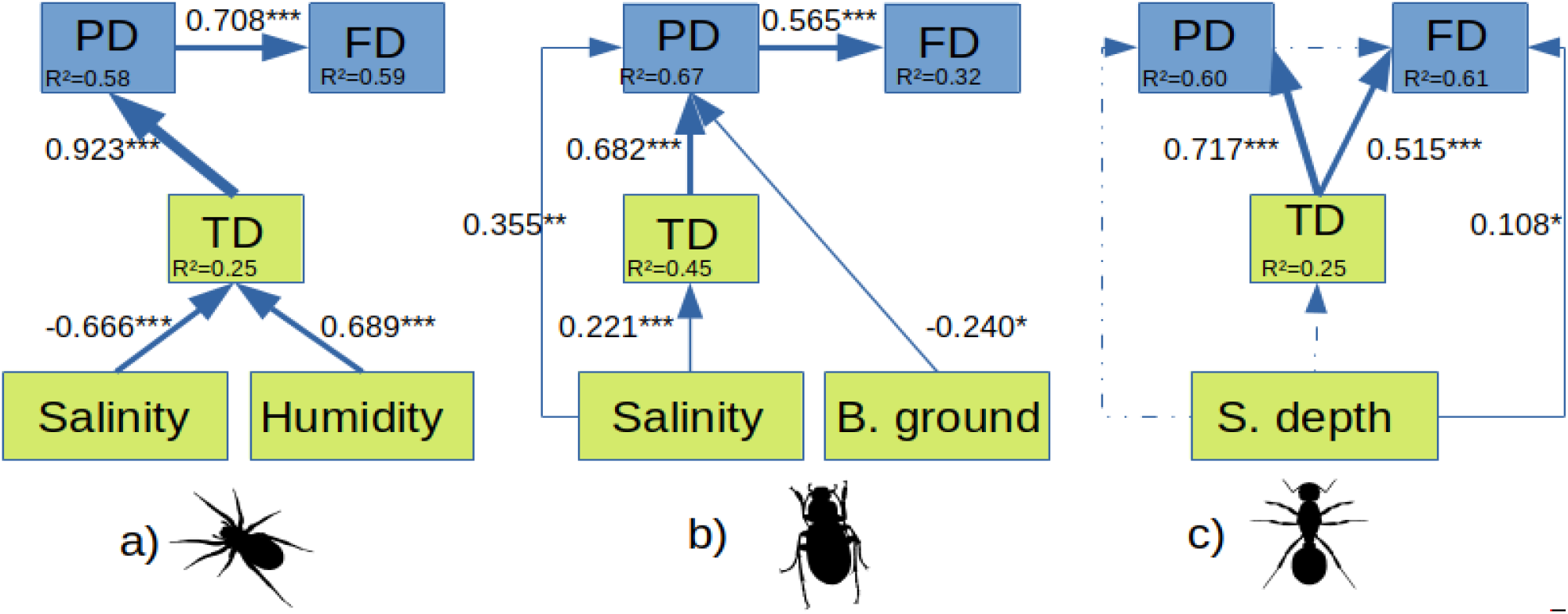
Best piecewise SEMs showing links between taxonomic, phylogenetic and functional diversity and environmental variables for a) spiders, b) carabid beetles and c) ants. Thickness of arrows is proportional to the standardized path coefficients (directionality and size given within boxes). Asterisks give significance level of linkages (<0.1, *<0.05, **<0.01, ***<0.001), and dashed lines correspond to paths included but not significant (p>0.05). Conditional R^2^ values are given within the boxes containing variables.

Environmental variables (here humidity and salinity indexes) were only linked to spider TD. Direction of these links were opposite with similar sizes. Spider phylogenetic diversity was strongly and positively related to TD while spider functional diversity was strongly and positively linked to phylogenetic diversity.

Similar (but weaker) relationships were found between carabid beetle α-, phylogenetic and functional diversities (Fig. 2). Environmental variables affecting carabid diversities were salinity and % bare-ground. Salinity positively influenced TD and phylogenetic diversity. % bare-ground had a negative effect on phylogenetic diversity.

Ant phylogenetic and functional diversities were strongly positively linked to TD. Soil depth was the only environmental variable selected by the model. It had a weak and positive effect on functional diversity. Selected, but not significant, paths were found between functional and phylogenetic diversities (p = 0.074), between soil depth and TD (p = 0.076) and between soil depth and phylogenetic diversity (p = 0.224).

## Discussion

Correlations were overall positive and high between FD and PD. These simple correlations may or may not represent mathematical artefacts from an increase of both, FD and PD with taxonomic diversity (TD). We will hence focus the discussion on the results that account for taxonomic diversity and other variables in parallel. Accounting for TD and environments made disappear this PD / FD relationship in ants, but maintained them in spiders and ground beetles, where TD related to FD only via increasing PD. Environmental constraints reduced FD in ants, but affected FD only indirectly via PD (ground beetles) and taxonomic and then PD (spiders and ground beetles).

### Differences of drivers between taxonomic, functional and phylogenetic diversities

The anthropogenic gradient of degradation, represented by bare ground, affected PD only in ground beetles, while this variable is known to affect TD of ground beetles in other habitats (see e.g. Pétillon et al. 2008 in salt marshes). Other environmental variables on the other hand have interacted with PD and FD in several ways, positively (humidity for spiders, salinity for ground beetles and soil depth for ants) or negatively (salinity for spiders), but most of their effects were indirect: With two exceptions (bare ground on PD of ground beetles and soil depth on FD of ants), environmental variables acted only on TD which then cascaded its effects onto PD which in turn affected FD. Environments affect taxonomic diversity rather than phylogenetic diversity, and taxonomic diversity affects phylogenetic diversity rather than functional diversity. Both, PD and FD are estimated using minimum spanning trees, but possibly a functional minimum spanning tree contains more polytomies, and hence adding a species without adding a new trait adds little branch length (see Appendix 2 for a simulation supporting this idea). Moreover, environments selecting for species may do so based on traits that we did not consider in the calculus of FD, and that are highly phylogenetically convergent, leading to a major increase in PD. Traits involved in stress response, like drought tolerance, indeed appear to have evolved convergently (Dunn et al. 1976). Increase in the diversity of the functional traits that we did consider would only be a side effect of such selection for phylogenetically convergent tolerances or resistances.

### Differences of drivers between taxa

Diversity patterns differing between taxa was one of our expectations (Wong et al. 2019) as assemblages of the three taxa are known to be differently affected by environmental variables depending on the ecology of species (e.g. Ridel et al. subm.). As an example, spiders diversity is less driven by landscape connectivity than ground beetles diversity (Lafage et al. 2015), and ants are strongly affected by local variables (Fichaux et al. 2019).

Spider TD responded negatively to salinity and drought. Salinity is a known environmental filter for spiders (Desender & Maelfait 1999; Pétillon et al. 2005; Traut 2005; Pétillon et al. 2007) hence the salinity resisting spider species are selected, reducing the local species richness. TD directly decreased with salinity, but PD and FD were only affected through TD. Salinity resistance, a trait not included in our analysis, thus may have appeared randomly in different parts of spider phylogenic tree (see Appendix 1): sorting for the few salinity tolerant species does not sort for phylogenetically particularly proximate or distant species. Humidity directly impacted only TD and impacted PD and FD only indirectly through TD. This could again be explained by environmental preferences and tolerances (not considered as traits here) if randomly distributed over spider phylogeny. Spider assemblage in the studied environment most likely result from a combination of humidity tolerant and mesohygrophilous species like in other ecotones (Traut 2005), which would explain the positive effect of humidity on spider TD (see also Entling et al. 2007, Weinninger & Fagan 2000 and Lafage et al. 2015; 2019). The positive effect of PD on FD in spiders suggests a phylogenetic signal of traits. This may appear contradictory with the tolerance traits considered above interpreted as being random across spider phylogeny, but these tolerance traits may not be representative of the whole spectrum of traits that characterize spiders. The majority of traits could show phylogenetic signal (Flynn et al. 2011; Khalil et al. 2018; Tucker et al. 2018). For instance, the hunting guild differs among families and hence shows strong phylogenetic signal (Cardoso et al. 2011).

TD and PD of ground beetles responded positively to salinity and negatively to bare ground. For ground beetles, contrary to spiders, salinity has a higher impact on PD than on TD. This would mean that salinity resistance is not restricted to a single or few phylogenetic branches but evolved convergently across distant branches of the phylogeny. Again, the absence of a direct effect of salinity on FD but only through PD can be explained by the spectrum of traits used here that may not affect salinity tolerance. The increase of TD with salinity appears inconsistent with existing literature from hypersaline habitats (Desender & Maelfait 1999; Pétillon et al. 2007). The high-cliff heathlands we studied are less saline and salinity might not exclude species, but only permit the additional occurrence of haloresistant species, therefore rising TD. The negative effect of bare ground on PD but not on TD suggests that bare ground selected for one, albeit species-rich phylogenetic lineage. Bare ground is anthropogenic here, and TD of ground beetles not declining under anthropogenic perturbation was previously reported (Verschoor & Krebs 1995), and possibly reflects quick recolonization post-perturbation by eurytopic and highly-dispersive species (Varet et al. 2013).

Ants were the only taxon in which FD was controlled by environmental constraints and TD. This was expected as FD is known to be very sensitive to environmental filters in ants (Campbell et al. 2010; Flynn et al. 2011, Arnan et al. 2015, Fichaux et al. 2019). A positive effect of soil depth on FD is consistent with the fact that most temperate species of ants nest underground (Torossian 1997) and hence profit from deep soils. Moreover, the increase of FD with TD, but not with PD suggests that traits accounted for in FD are not phylogenetically conserved nor convergent in ants. Instead, adding a species seems to on average add trait values to the community, independent of the selection pressure of environmental constraints.

## Conclusions

Overall, among the models formulated in the Introduction, results are consistent with phylogenetic conservatism in traits in spiders and ground beetles: phylogenetic diversity relates positively to functional diversity (Webb et al. 2002), even when accounting for taxonomic diversity. For ants, in contrast, results appear to be more consistent with a more neutral model with any new species potentially representing a new trait state (Purves & Turnbull 2010, Novack-Gottshall 2016; Stevens & Grimshaw 2020). The fact that phylogenetic diversity often increases mainly with taxonomic diversities, might reflect either an effect of an unmeasured environmental condition (e.g. weather condition as they are known to be harsh in the studied habitat see Sawtschuck 2010) affecting both taxonomic diversity and phylogenetic diversity, or a coexistence facilitated among phylogeneticaly distant species, resulting in a positive relationship between taxonomic and phylogenetic diversity. Our results are not consistent with models of character displacement among close relatives that should lead to negative relationships between phylogenetic and functional diversity (Prinzing et al. 2008), or reduced niche packing under environmental constraints (Safi et al. 2011). These models invoke competitive interactions that might be low in some ground-dwelling predators (see Wise 2006 for spiders and Fichaux et al. 2019 for ants).

## Acknowledgments

We thank Clément Gouraud for his expertise regarding ants, Aurélien Ridel, Pierre Devogel and Timothée Scherer for their help in identifying spiders and carabids, and sorting samples respectively, and “Bretagne Vivante” and “Communauté de communes de Belle-île-en-Mer” for continuous support during fieldwork.

## Declarations

### Funding

This study was funded by the Fondation de France (grant number n° 1531).

### Conflicts of interest

The authors declare that they have no conflict of interest.

### Ethic approval

This article does not contain any studies with human participants or animals performed by any of the authors.

### Consent to participate (include appropriate statements)

all authors

### Consent for publication (include appropriate statements)

all authors

### Availability of data and material (data transparency)

Dryad?

### Code availability (software application or custom code)

Not applicable

---

Appendix 1: phylogenetic tree of spider species sampled in this study. The colors marks depend on each specie’s salt tolerance (Green= Halophilic; bleu= tolerant; red= intolerant; no mark= NA).

Appendix 2: Functional diversity simulation

## Notes

### Competing Interest Statement

The authors have declared no competing interest.

## Bibliography

Agnarsson I (2004) Morphological phylogeny of cobweb spiders and their relatives (Araneae, Araneoidea, Theridiidae). Zool J Linn Soc 141(4):447–626 doi: 10.1111/j.1096-3642.2004.00120.x

Arnan X, Cerdá X, Retana J (2015) Partitioning the impact of environment and spatial structure on alpha and beta components of taxonomic, functional, and phylogenetic diversity in European ants. PeerJ 3:e1241 doi: 10.7717/peerj.1241

Arnedo MA, Hormiga G, Scharff N (2009) Higher-level phylogenetics of linyphiid spiders (Araneae, Linyphiidae) based on morphological and molecular evidence. Cladistics 25(3):231–262 doi: 10.1111/j.1096-0031.2009.00249.x

Azevedo GHF, Griswold CE, Santos, AJ (2018) Systematics and evolution of ground spiders revisited (Araneae, Dionycha, Gnaphosidae). Cladistics 34(6):579–626 doi: 10.1111/cla.12226

Bernard-Verdier M, Flores O, Navas ML, Garnier E. (2013) Partitioning phylogenetic and functional diversity into alpha and beta components along an environmental gradient in a Mediterranean rangeland. J Veg Sci 24(5):877–889

Blatrix R, Galkowski C, Lebas C, Wegnez P (2013) Guide des fourmis de France, Delachaux et Niestlé

Bond JE, Garrison NL, Hamilton CA, Godwin RL, Hedin M, Agnarsson I (2014) Phylogenomics resolves a spider backbone phylogeny and rejects a prevailing paradigm for orb web evolution. Curr Biol 24(15):1765–1771

Bürkner P-C (2018) Advanced bayesian multilevel modeling with the R package brms. R J 10(1):395–411 doi: 10.32614/RJ-2018-017

Cadotte MW, Carboni M, Si X, Tatsumi S (2019) Do traits and phylogeny support congruent community diversity patterns and assembly inferences?. J Ecol 107(5):2065–2077

Campbell WB, Freeman DC, Emlen JM, Ortiz SL (2010) Correlations between plant phylogenetic and functional diversity in a high altitude cold salt desert depend on sheep grazing season: Implications for range recovery. Ecol Indic 10(3):676–686

Cardoso P, Pekár S, Jocqué R, Coddington JA (2011) Global patterns of guild composition and functional diversity of spiders. PloS one 6(6)

Cardoso P, Rigal F, Carvalho JC (2015) BAT–Biodiversity assessment tools, an R package for the measurement and estimation of alpha and beta taxon, phylogenetic and functional diversity. Methods Ecol Evol 6(2):232–236

Cavender-Bares J, Kozak KH, Fine PV, Kembel SW (2009) The merging of community ecology and phylogenetic biology. Ecol Lett 12(7):693–715

Coelho MS, Quintino AV, Fernandes GW, Santos JC, Delabie JHC (2009) Ants Hymenoptera: Formicidae) as bioindicators of land restoration in a Brazilian Atlantic forest fragment. Sociobiology 54(1):51

Corbelli JM, Zurita GA, Filloy J, Galvis JP, Vespa NI, Bellocq I (2015) Integrating taxonomic, functional and phylogenetic beta diversities: Interactive effects with the biome and land use across taxa. PloS one 10(5)

Dunn EL, Shropshire FM, Song LC, Mooney HA (1976) The water factor and convergent evolution in Mediterranean-type vegetation. In: Water and plant life. Springer, Berlin, Heidelberg, pp 492–505

Ellenberg H, Weber HE, Düll R, et al (1992) Zeigerwerte von Pflanzen in Mitteleuropa, 2nd ed. Scr Geobot 18:1–258

Entling W, Schmidt MH, Bacher S, Brandl R, Nentwig W (2007) Niche properties of Central European spiders: shading, moisture and the evolution of the habitat niche. Glob Ecol Biogeogr, 16(4):440–448

Eterovick PC, Rievers CR, Kopp K, Wachlevski M, Franco BP, Dias CJ, Barata IM, Ferreira ADM, Afonso LG (2010) Lack of phylogenetic signal in the variation in anuran microhabitat use in southeastern Brazil. Evol Ecol 1–24

Faith DP (1992) Conservation evaluation and phylogenetic diversity. Biol Conserv 45761:1–10

Fichaux M, Béchade B, Donald J, Weyna A, Delabie JHC, Murienne J, Baraloto C, Orivel J (2019) Habitats shape taxonomic and functional composition of Neotropical ant assemblages. Oecologia 189(2):501–513

Flynn DF, Mirotchnick N, Jain M, Palmer MI, Naeem S (2011) Functional and phylogenetic diversity as predictors of biodiversity–ecosystem-function relationships. Ecology 92(8):1573–1581

Frick H, Nentwig W, Kropf C (2010) Progress in erigonine spider phylogeny—the Savignia-group is not monophyletic (Araneae?: Linyphiidae). Org Divers Evol 10(4):297–310 doi: 10.1007/s13127-010-0023-1

Gerlach J, Samways M, Pryke J (2013) Terrestrial invertebrates as bioindicators: an overview of available taxonomic groups. J Insect Conserv 17(4):831–850 doi: 0.1007/s10841-013-9565-9

Grundler MR, Pianka ER, Pelegrin N, Cowan MA, Rabosky DL (2018) Stable isotope ecology of a hyper-diverse community of scincid lizards from arid Australia. PLOS One 12: e0172879

Haak, DC, Ballenger BA, Moyle LC (2014) No evidence for phylogenetic constraint on natural defense evolution among wild tomatoes. Ecology 95:1633–1641

Hacala A, Le Roy M, Sawtschuk J, Pétillon J (2020) Comparative responses of spiders and plants to maritime heathland restoration. Biodivers Conserv 29(1):229–249

Hill MO, Preston CD, Roy DB (2004) PLANTATT-attributes of British and Irish plants: status, size, life history, geography and habitats. Centre for Ecology & Hydrology.

Jeannel R (1941) Faune de France: Coléoptères carabiques. Fédération Francaise des sociétés de science naturelles. Office central de faunistique.

Khalil MI, Gibson DJ, Baer SG, Willand JE (2018) Functional diversity is more sensitive to biotic filters than phylogenetic diversity during community assembly. Ecosphere, 9(3):e02164. 10.1002/ecs2.2164

Kruschke JK, Liddell TM (2018) The Bayesian New Statistics: Hypothesis testing, estimation, meta-analysis, and power analysis from a Bayesian perspective. Psychon Bull Rev, 25(1):178–206

Lafage D, Maugenest S, Bouzillé J-B, Pétillon J (2015) Disentangling the influence of local and landscape factors on alpha and beta diversities: opposite response of plants and ground-dwelling arthropods in wet meadows. Ecol Res 30:1025–1035

Lafage D, Djoudi EA, Perrin G, Gallet S, Pétillon J (2019) Responses of ground-dwelling spider assemblages to changes in vegetation from wet oligotrophic habitats of Western France. Arthropod Plant Interact 13:653–662

Laliberté E, Legendre P, Shipley B, Laliberté ME (2014) Package ‘FD’. Measuring functional diversity from multiple traits, and other tools for functional ecology

Lefcheck JS (2016) piecewiseSEM: Piecewise structural equation modelling in r for ecology, evolution, and systematics. Methods Ecol Evol, 7(5):573–579

Leroy B, Le Viol I, Pétillon J (2014) Complementarity of rarity, specialisation and functional diversity metrics to assess community responses to environmental changes, using an example of spider communities in salt marshes. Ecol Indic 46:351–357

Liu C, Guénard B, Blanchard B, Peng YQ, Economo EP (2016) Reorganization of taxonomic, functional, and phylogenetic ant biodiversity after conversion to rubber plantation. Ecol Monogr 86(2):215–227 doi: 10.1890/15-1464.1

López-López A, Vogler AP (2017) The mitogenome phylogeny of Adephaga (Coleoptera). Mol Phylogenet Evol 114:166–174 doi: 10.1016/j.ympev.2017.06.009

Luff ML (2007) The Carabidae (ground beetles) of Britain and Ireland. RES handbooks for the identification of British Insects Vol 4 Part 2. Field Studies Council, Shrewsbury

Maddison WP (2015) A phylogenetic classification of jumping spiders (Araneae?: Salticidae). J Arachnol 43(3):231 doi: 10.1636/arac-43-03-231-292

Magura T (2016) Ignoring functional and phylogenetic features masks the edge influence on ground beetle diversity across forest-grassland gradient. For Ecol Manage 384:371–377 doi: 10.1016/j.foreco.2016.10.056

Makowski D, Ben-Shachar MS, Lüdecke D (2019) bayestestR: Describing effects and their uncertainty, existence and significance within the Bayesian framework. J Open Source Softw, 4(40):1541

Martínez-Navarro EM, Galián J, Serrano J (2005) Phylogeny and molecular evolution of the tribe Harpalini (Coleoptera, Carabidae) inferred from mitochondrial cytochrome-oxidase I. Mol Phylogenetics Evol 35(1):127–146 doi:10.1016/j.ympev.2004.11.009

Millidge AF (1977) The genera Mecopisthes Simon and Hypso-cephalus n.gen. and their phylogenetic relationships Araneae:Linyphiidae. Bulletin of the British Arachnological Society(4):113–123

Moreau CS, Bell CD, Vila R, Archibald SB, Pierce NE (2006) Phylogeny of the Ants?: Diversification in the Age of Angiosperms. Science, 312(5770):101–104 doi: 10.1126/science.1124891

Nentwig W, Blick T, Gloor D, Hänggi A, Kropf C (2019) Version 06.2019. Online at https://www.araneae.nmbe.ch. Accessed 19 June 2019. doi: 10.24436/1

Novack-Gottshall PM (2016) General models of ecological diversification. I. Conceptual synthesis. Paleobiology 42:185–208

Ober KA, Maddison DR (2008) Phylogenetic relationships of tribes within Harpalinae (Coleoptera?: Carabidae) as inferred from 28S ribosomal DNA and the wingless gene. J Insect Sci 8(63):1–32 doi:10.1673/031.008.6301

Paradis E, Blomberg S, Bolker B, Brown J, Claude J, Cuong HS, Desper R (2019) Package ‘ape’. Analyses of phylogenetics and evolution, version, 2(4)

Pavoine S, Baguette M, Bonsall MB (2010) Decomposition of trait diversity among the nodes of a phylogenetic tree. Ecol Monogr, 80(3):485–507

Pavoine S, Gasc A, Bonsall MB, Mason NW (2013) Correlations between phylogenetic and functional diversity: mathematical artefacts or true ecological and evolutionary processes?. J Veg Sci, 24(5):781–793.

Parr CL, Dunn RR, Sanders NJ, Weiser MD, Photakis M, Bishop TR, Fitzpatrick MC, Arnan X, Baccaro F, Brand∼Ao CRF, Chick L, Donoso DA, Fayle TM, Gomez C, Munyai BTC, Pacheco R, Retana J, Robinson A, Sagata K, Silva RR, Tista M, Vasconcelos H, Yates M, Gibb H (2017) GlobalAnts: A new database on the geography of ant traits (Hymenoptera: Formicidae). Insect Conserv Divers 10(1):5–20

Pedley SM, Dolman PM (2014) Multi-taxa trait and functional responses to physical disturbance. J Anim Ecol 83(6):1542–1552 doi: 10.1111/1365-2656.12249

Pétillon J, Ysnel F, Canard A, Lefeuvre J-C (2005) Impact of an invasive plant (Elymus athericus) on the conservation value of tidal salt marshes in western France and implications for management: responses of spider populations. Biol Conserv 126:103–117

Pétillon J, Georges A, Canard A, Ysnel F (2007) Impact of cutting and sheep grazing on ground–active spiders and carabids in intertidal salt marshes (Western France). Anim Biodivers Conserv, 30(2):201–209

Pétillon J, Georges A, Canard A, Lefeuvre J-C, Bakker JP, Ysnel F (2008) Influence of abiotic factors on spider and ground beetles communities in different salt-marsh systems. Basic and Applied Ecology 9:743–751

Pétillon J, Potier S, Carpentier A, Garbutt A (2014) Evaluating the success of managed realignment for the restoration of salt marshes: Lessons from invertebrate communities. Ecol Eng 69:70–75

Piacentini LN, Ramírez MJ (2019) Hunting the wolf?: A molecular phylogeny of the wolf spiders (Araneae, Lycosidae). Mol Phylogenet Evol 136:227–240

Pinheiro-Silva L, Gianuca AT, Silveira MH, Petrucio MM (2020) Grazing efficiency asymmetry drives zooplankton top-down control on phytoplankton in a subtropical lake dominated by non-toxic cyanobacteria. Hydrobiologia, 1-14

Prinzing A, Reiffers R, Braakhekke WG, Hennekens SM, Tackenberg O, Ozinga WA, Schaminée JHJ, Van Groenendael JM (2008) Less lineages–more trait variation: phylogenetically clustered plant communities are functionally more diverse. Ecol Lett, 11(8), 809-819.

Purves DW, Turnbull LA (2010) Different but equal: the implausible assumption at the heart of neutral theory. J Anim Ecol 79:1215–1225

Ridel, A., Lafage, D., Devogel, P., Lacoue-Labarthe, T., Pétillon, J. (2020) Habitat filtering differentially modulates phylogenetic vs functional diversity relationships between dominant ground-dwelling arthropods in salt marshes. DOI: https://doi.org/10.1101/2020.06.19.161588

Roberts MJ (1985) The spiders of Great Britain and Ireland. Brill Archive

Rodrigues ASL, Brooks TM, Gaston KJ (2005) Integrating phylogenetic diversity in the selection of priority areas for conservation: does it make a difference. Phylogeny and conservation, 8:101–119.

Ruiz C, Jordal B, Serrano J (2009) Molecular phylogeny of the tribe Sphodrini (Coleoptera?: Carabidae) based on mitochondrial and nuclear markers. Mol Phylogenet Evol 50(1):44–58 doi: 10.1016/j.ympev.2008.09.023

Sasakawa K, Kubota K (2007) Phylogeny and genital evolution of Carabid beetles in the genus Pterostichus and its allied genera (Coleoptera?: Carabidae) inferred from two nuclear gene sequences. Ann Entomol Soc Am 100(2):100–109 doi: 10.1603/0013-8746(2007)100[100:PAGEOC]2.0.CO;2

Safi K, Cianciaruso MV, Loyola RD, Brito D, Armour-Marshall K, Diniz-Filho JAF (2011) Understanding global patterns of mammalian functional and phylogenetic diversity. Philos Trans R Soc Lond B Biol Sci 366(1577):2536–2544

Scharff N, Coddington JA, Blackledge TA, Agnarsson I, Framenau VW, Szűts T, Hayashi CY, Dimitrov D (2019) Phylogeny of the orb-weaving spider family Araneidae (Araneae?: Araneoidea). Cladistics cla.12382 doi: 10.1111/cla.12382

Sawtschuk J (2010) Restauration écologique des pelouses et des landes des falaises littorals atlantiques: Analyse des trajectoires successionnelles en environnement contraint. Phd dissertation, Institut de Géoarchitecture, Université de Bretagne Occidentale, Brest, France.

Schirmel J, Blindow I, Buchholz S (2012) Life-history trait and functional diversity patterns of ground beetles and spiders along a coastal heathland successional gradient. Basic and Applied Ecology 13:606–614 doi: 10.1016/j.baae.2012.08.015

Schirmel J, Thiele J, Entling MH, Buchholz S (2016) Trait composition and functional diversity of spiders and carabids in linear landscape elements. Agric Ecosyst Environ 235:318–328 doi: 10.1016/j.agee.2016.10.028

Stevens RD, Grimshaw JR (2020) Relative contributions of ecological drift and selection on bat community structure in interior Atlantic Forest of Paraguay. Oecologia 193: 645–654

Topping CJ, Sunderland KD (1992) Limitations to the use of pitfall traps in ecological studies exemplified by a study of spiders in a field of winter wheat. J Appl Ecol 29(2):485–491

Torossian C (1977) Les fourmis rousses des bois (Formica rufa) indicateurs biologiques de dégradation des forêts de montagne des Pyrénées orientales. Bull d’Écol 8(3):333–348

Traut BH (2005) The role of coastal ecotones: a case study of the salt marsh/upland transition zone in California. J Ecol 279-290

Tucker CM, Davies TJ, Cadotte MW, Pearse WD (2018) On the relationship between phylogenetic diversity and trait diversity. Ecology 99(6):1473–1479

Varet M, Burel F, Lafage D, Pétillon J (2013) Age-dependent colonization of urban habitats: a diachronic approach using carabid beetles and spiders. Anim Biol 63:257–269

Verschoor BC, Krebs BPM (1995) Diversity changes in a plant and carabid community during early succession in an embanked saltmarsh area. Pedobiologia 39:405–416.

Violle C, Navas ML, Vile D, Kazakou E, Fortunel C, Hummel I, Garnier E (2007) Let the concept of trait be functional! Oikos 116(5):882–892

Wang F, Ballesteros JA, Hormiga G, Chesters D, Zhan Y, Sun N, Zhu C, Chen W, Tu L (2015) Resolving the phylogeny of a speciose spider group, the family Linyphiidae (Araneae). Mol Phylogenet Evol 91:135–149 doi: 10.1016/j.ympev.2015.05.005

Webb CO, Ackerly DD, McPeek MA, Donoghue MJ (2002) Phylogenies and community ecology. Annu Rev Ecol Syst 33(1):475–505

Wenninger EJ, Fagan WF (2000) Effect of river flow manipulation on wolf spider assemblages at three desert riparian sites. J Arachnol, 28(1):115–122

Wheeler WC, Coddington JA, Crowley LM, Dimitrov D, Goloboff PA, Griswold CE, Hormiga G, Prendini L, Almeida-Silva L, Alvarez-Padilla F, Arnedo MA, Benavides Silva LR, Benjamin SP, Bond JE, Grismado CJ, Hasan E, Hedin M, Izquierdo MA, Labarque FM, Ledford J, Lopardo L, Maddison WP, Miller JA, Piacentini LN, Platnik NI, Polotow D, Silva-Dávila D, Scharff N, Szűts T, Ubick D, Vink CJ, Wood HM, Zhang, J (2017) The spider tree of life?: Phylogeny of Araneae based on target-gene analyses from an extensive taxon sampling. Cladistics 33(6):574–616 doi: 10.1111/cla.12182

Wiens JJ, Ackerly DD, Allen AP, Anacker BL, Buckley LB, Cornell HV, Damschen EI, Davis JT, Grytnes J-A, Harrison SP, Hawkins BA, Holt RD, McCain CM, Stephens PR (2010) Niche conservatism as an emerging principle in ecology and conservation biology. Ecol lett, 13(10):1310–1324

Wise DH (2006) Cannibalism, food limitation, intraspecific competition and the regulation of spider populations. Annu Rev Entomol 51:441–465

Wong MK, Guénard B, Lewis OT (2019) Trait-based ecology of terrestrial arthropods. Biol Rev 94(3):999–1022

