## Supplementary figures and images for "Drivers of taxonomic, functional and phylogenetic diversities in dominant ground-dwelling arthropods of coastal heathlands"

### appendix 1

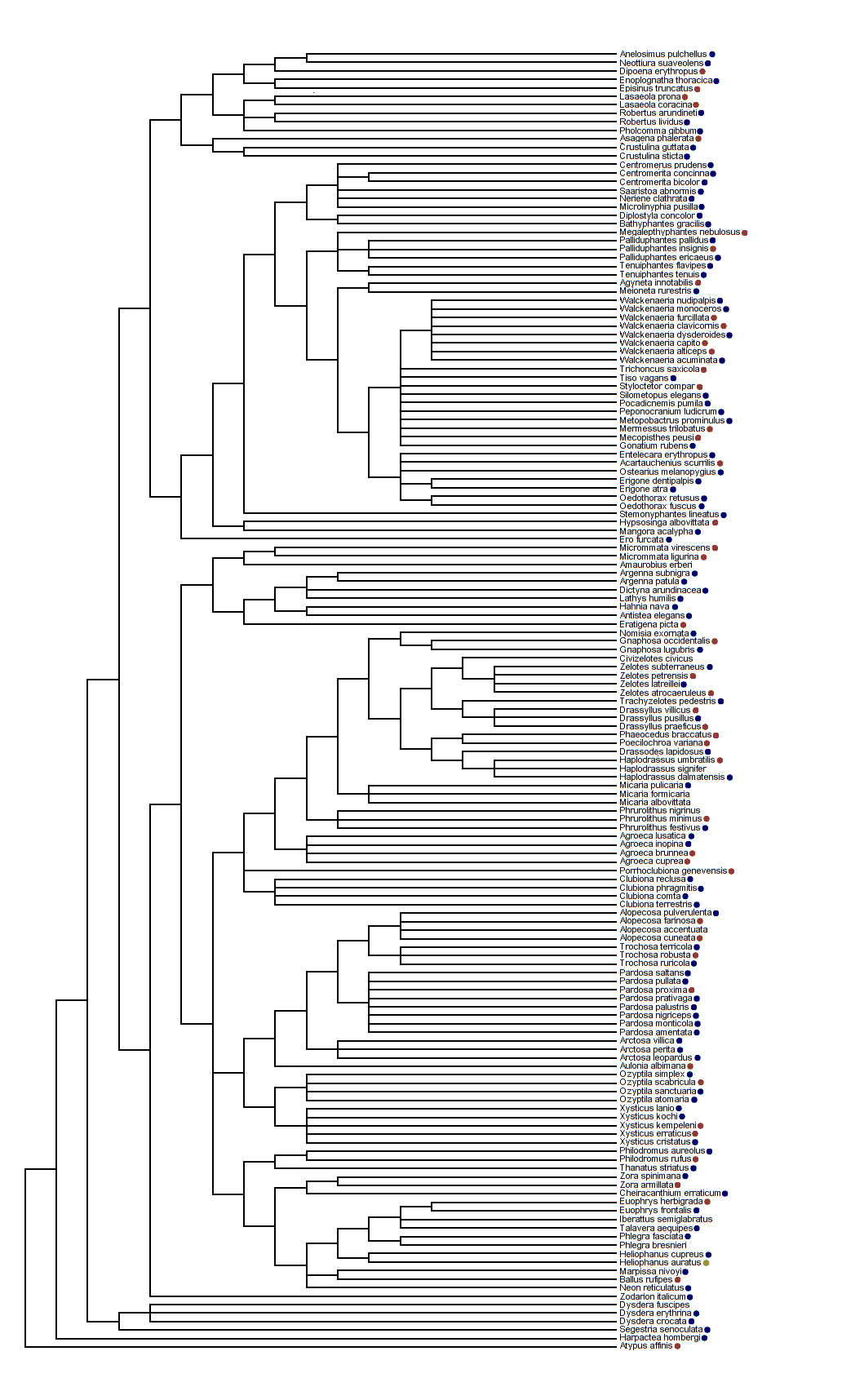
