## appendix 2 for "Drivers of taxonomic, functional and phylogenetic diversities in dominant ground-dwelling arthropods of coastal heathlands"

Method

In order to simulate the effect on functional diversity of adding species with identical traits in a community, we added 10 simulated species with traits similar to Haplodrassus_dalmatensis to each community sample. We then calculated the FD value using the BAT package for the simulated and real communities. Distance matrices were computed with gower distance from the FD package (Laliberté et al. 2014). Mean FD values for all the samples were then compared using a Wilcoxon test.

Results

| Site | Degradation state | Mean FD for simulated communities | Mean FD for real communities |
| --- | --- | --- | --- |
| PH | RF | 7.31 | 8.61 |
| PH | HD | 3.01 | 3.07 |
| ENF | HD | 4.35 | 4.45 |
| PH | MD | 4.05 | 4.64 |
| ENF | HD | 5.01 | 5.03 |
| ENF | HD | 3.70 | 3.79 |
| ENF | MD | 3.46 | 3.59 |
| ENF | MD | 7.51 | 7.63 |
| ENF | MD | 8.66 | 8.82 |
| ENF | MD | 2.98 | 3.08 |
| ENF | MD | 7.15 | 6.98 |
| APO | HD | 5.81 | 5.94 |
| ENF | MD | 4.12 | 4.24 |
| APO | HD | 6.57 | 6.70 |
| ENF | HD | 5.82 | 5.89 |
| APO | HD | 5.38 | 5.54 |
| APO | MD | 5.20 | 5.31 |
| APO | RF | 6.29 | 6.38 |
| ENF | HD | 7.94 | 7.97 |
| ENF | HD | 5.36 | 5.41 |
| PH | MD | 6.23 | 6.23 |
| PH | HD | 5.74 | 5.72 |
| PH | HD | 5.18 | 5.22 |
| PH | MD | 7.55 | 7.62 |
| APO | HD | 6.54 | 6.57 |
| APO | HD | 6.17 | 6.30 |
| PH | RF | 4.84 | 4.88 |
| PH | HD | 5.48 | 5.53 |
| PH | HD | 3.01 | 3.12 |
| ENF | MD | 7.34 | 7.36 |
| PH | MD | 6.09 | 6.23 |
| PH | MD | 7.14 | 7.15 |
| PH | MD | 5.52 | 5.47 |

The mean FD value for the simulated communities was not significantly different from the mean FD value for the real communities (W = 513, P = 0.693).
